# Longitudinal analysis of drift in the circulating human antibody repertoire over four years

**DOI:** 10.1101/2025.03.20.644446

**Authors:** Collin Joyce, Benjamin Nemoz, Raiza Bastidas, Bryan Briney, Dennis R. Burton

**Affiliations:** Department of Immunology and Microbiology, The Scripps Research Institute, La Jolla, CA, USA; Scripps Consortium for HIV/AIDS Vaccine Development, The Scripps Research Institute, La Jolla, CA, USA; IAVI Neutralizing Antibody Center, The Scripps Research Institute, La Jolla, CA, USA; Center for Viral Systems Biology, The Scripps Research Institute, La Jolla, CA, USA; Multi-Omics Vaccine Evaluation Consortium, The Scripps Research Institute, La Jolla, CA, USA; Ragon Institute of MGH, MIT and Harvard, Cambridge, MA, USA

## Abstract

The human immune system is a sophisticated network of cells and molecules that plays a vital role in safeguarding the body against a broad range of foreign agents. One of the key components of the immune system is the circulating antibody repertoire, a vast collection of different antibodies that recognize and eliminate foreign agents specifically. The diversity and specificity of the circulating antibody repertoire are essential for effective immunity. In a study of ten healthy donors, we previously demonstrated the accessible human antibody repertoire to be on the order of 10^15^-10^18^ distinct members, although an individual will only sample a fraction of that repertoire at a given time point. The mode of generation of the naïve antibody repertoire through random recombination before antigen contact suggests that the composition of the circulating antibody repertoire may drift over time, but the magnitude of this drift is poorly understood. Here, we used high-throughput sequencing to analyze two donors from our original study, approximately four years later. Our results reveal conservation of the size, overall diversity, and gross features of the circulating antibody repertoire, such as V- and J-gene usage and CDRH3 length distribution, over time but suggest substantial changes to fine features of the repertoire, such as combinations of V, J, D gene segments and P and N sequences, which may influence primary responses to newly encountered pathogens as well as to immunization procedures.

## INTRODUCTION

Antibodies can theoretically be raised to bind to virtually any non-self-antigen. This impressive capability is associated with a vast diverse repertoire of antibodies that is initiated during B cell development in the bone marrow. Three unlinked loci contain the gene segments that make up an antibody: the heavy chain locus on chromosome 14 and two light chain loci on chromosomes 2 and 22 (lambda and kappa chains, respectively). The heavy chain variable domain is assembled from three gene segments VH, DH, and JH, that are randomly joined. In humans, there are ∼39-56 VH, 23-27 DH and 6 JH gene segments (the precise numbers differ between individuals) to generate ∼ 10^4^ different combinations. Recombination requires V(D)J recombinase, a protein complex that includes RAG1, RAG2 and Artemis. Artemis and TdT are responsible for the random (non-templated) addition of P and N nucleotides at VH-DH and DH-JH junctions, dramatically increasing sequence diversity. This newly formed junctional region that contains the C-terminal end of the VH gene segment, the entire DH gene segment, and the N-terminal beginning of the JH gene segment corresponds to the heavy chain complementarity determining region 3 (CDRH3), which is typically crucial in antigen recognition. Following the successful pairing of the newly formed heavy chain with the surrogate light chain (SLC), the B cell undergoes recombination of a light chain from the V and J gene segments of either the kappa or lambda loci. In humans, there are ∼300-400 combinations generated theoretically by VJ recombination that can be expanded by N addition. The SLC is then replaced with this newly generated light chain. If the immature B cell is not autoreactive or anergic and thus does not undergo receptor editing or clonal deletion, it matures into a naïve B cell. This mature B cell migrates to the periphery, where it can be activated upon encountering an antigen where diversity is further considerably amplified by somatic hypermutation^1^. From the above, the naïve antibody repertoire is of the order of 10^6^ that is greatly amplified by N and P addition. Indeed, comprehensive analysis of human antibody repertoires has historically been hampered by their immense size^2,3^. Additionally, from a practical standpoint, it should be noted that a substantial portion of the B lymphocyte population is sequestered in organs and tissues that are not readily accessible for sampling in living subjects so that studies of human repertoires have predominantly centered on circulating B cells^4^. Furthermore, this approach inherently has limitations, as only a subset of circulating B cells can be sampled at any given time.

We previously examined the circulating B cell populations of ten human subjects and described the sequencing of almost 3 billion antibody heavy chain sequences, which constituted, at the time (2019), the largest single collection of adaptive immune receptor sequences^5^. We concluded that the accessible human naïve antibody repertoire was orders of magnitude larger than had been previously estimated and was in the range of 10^15^-10^18^ unique antibody molecules. The study further revealed a largely unique repertoire for each of the individuals investigated and a subpopulation of universally shared “clonotypes”, denoted “public clonotypes”. An antibody clonotype is here defined as a collection of sequences using the same variable (V) and joining (J) genes and encoding an identical CDRH3 amino acid sequence^5,6^ and allows minimization of the effects of sequencing and amplification errors, while also rigorously controlling for clonal lineage size.

Prior studies have suggested that key features of the circulating antibody repertoire, such as V-gene and J-gene usage, CDRH3 length distributions and clonotypic composition may drift over time within a single individual, but the magnitude of this drift is poorly understood due to limited sampling depth^2,3^ or a restriction to specific cell types^6-9^. To further our understanding of how the composition of these repertoire features changes over time, we re-sequenced two of the ten previously studied subjects after a four-year interval and compared these new data with the initial timepoint.

## RESULTS

The two donors who were available for re-sequencing were D103 and 327059^5^ (Table S1). We followed the same protocols as previously described. Eighteen new libraries per subject yielded 6.2 × 10^8^ raw reads. After annotation^10^, filtering, and duplicate removal using unique molecular identifiers^11^, we obtained a total of 6.1 × 10^7^ productive antibody sequences across both donors for this new timepoint.

### High-level genetic repertoire feature comparisons between timepoints for each donor

First, we investigated changes to high-level genetic repertoire features, namely V-gene and J-gene usage and CDRH3 length distributions. Consistent with prior observation^2,6-9^, overall VH gene usage was largely stable over time, although VH gene use was somewhat less correlated across timepoints than between biological replicates at the same time point (Figure 1a, Figures S1-2). The frequencies of IgM-encoding (0.76–0.94) and IgG-encoding (0.06–0.24) sequences were consistent with expected frequencies^12^ (Figure 1b). Similar V-gene, J-gene and CDRH3 length distributions were observed (Figure 1c, e, f) at each time point. We aimed to identify a set of features that could be used to classify or timestamp repertoires that was more granular than any one feature alone. For this, we settled on collapsing repertoires into sets of V gene, J gene and CDRH3 length combinations and quantified similarity using the Morisita–Horn similarity index^13,14^ (refer to Materials and Methods). This combination of features was designed to emulate the level of granularity achieved by using *species* as the quantifiable unit in ecological similarity analyses. The use of individual genomes (analogous to individual antibody sequences) to compute ecological similarity is not feasible, since the frequency of identical twins is sufficiently low that analyses premised on the repeat occurrence of identical genomes in two populations are not informative. Instead, these methods require the use of a higher-order classification, like species in the ecological case. In our case, donor-matched samples from different timepoints were clearly distinguishable using as few as 10^4^ sequences (Figure 1d) with intra-timepoint and donor biological replicates clustered together (Figure 1g). Additionally, Figure 1g shows neither inter-donor nor intra-donor but inter-timepoint biological replicates clustered together.

**Figure 1.**
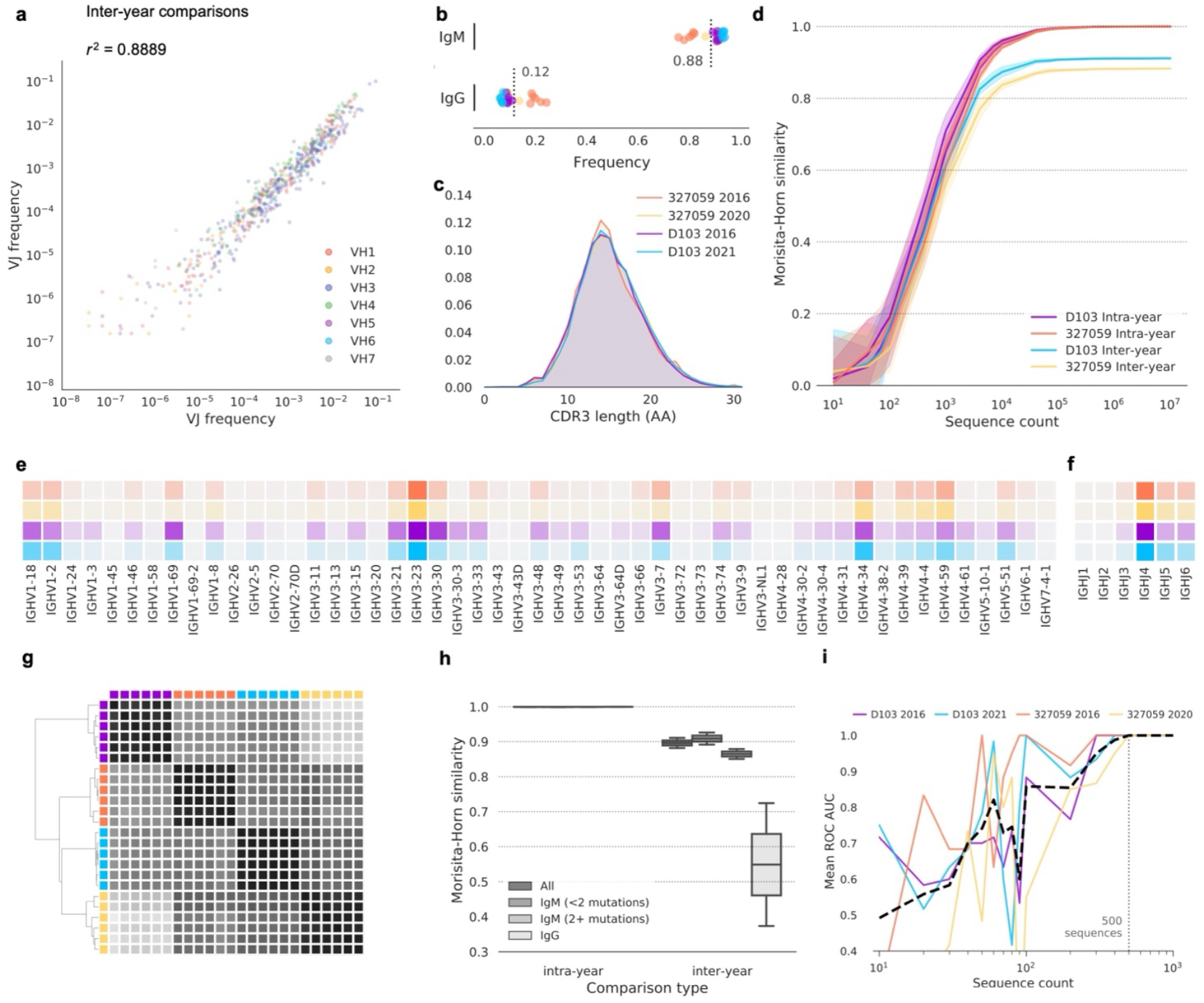
Uniqueness of the repertoires at individual timepoints. (a) Frequency comparison of V and J combinations between timepoints. V and J combinations are colored according to the V gene used. (b) Sequence frequency by antibody isotype. Timepoints are colored as in c. Each point represents a single biological replicate. Mean of all samples is indicated for each isotype. (c) CDRH3 length distribution for each timepoint. CDRH3 lengths were determined using the Immunogenetics (IMGT) numbering scheme. AA, amino acids. TP, timepoint. (d) Intra- and inter-year pairwise Morisita–Horn similarity comparisons for each subject. Lines indicate mean similarity of 20 bootstrap samplings, and shaded areas indicate 95% confidence intervals. V gene (e) and J gene (f) use by subject. Increased color intensity indicates higher frequency. Subjects are colored as in (c). (g) Clustered distance matrix of biological replicates from each subject and timepoint, using pairwise Morisita–Horn similarity of V-gene, J-gene and CDRH3 length as the distance measure. Distance matrix was computed using single-linkage clustering (Euclidean distance metric). Timepoint colors are as in (c). A dendrogram representation of the distance matrix is also shown. (h) Comparison of intra- and inter-timepoint similarity in V-gene, J-gene and CDRH3 length, using all sequences, IgM sequences with fewer than two nucleotide mutations, IgM sequences with two or more mutations, or IgG sequences. Box plots show the median line and span the 25th–75th percentile, with whiskers indicating the 95% confidence interval. (j) Mean receiver operating characteristic (ROC) area under the curve (AUC) for a one-versus-rest support-vector-machine classifier. The ROC AUC does not drop below 1.0 for any subject when the test or training datasets include ≥ 500 sequences; this 500-sequence threshold is indicated with a dashed vertical line.

IgG+ repertoires were less similar than IgM repertoires (Figure 1h), suggesting new immunological encounters impact the conservation of high-level features more than ongoing naive B cell replacement from the bone marrow. A one-versus-rest support-vector-machine classifier trained on V-gene, J-gene and CDRH3 length data from 5 of the 6 biological replicates from each sampled repertoire accurately assigned the remaining replicate using test or training datasets of as few as 500 sequences from each replicate (Figure 1i), indicating a significant change in the circulating repertoire over time which was large enough to be observed even when distilling the repertoire to a frequency distribution of only three genetic features.

### Calculation and characterization of sequence and clonotype persistence between timepoints for each donor

For each subject, all sequences from each timepoint were collapsed into a set of unique clonotypes. The fraction of clonotypes and nucleotide sequences shared between samples from different timepoints for a given donor was found to increase linearly with sampling depth (Figure S3). A total of 32,834 and 167,941 clonotypes for subjects D103 and 327059 respectively were shared across timepoints (Figure 2a-b). To estimate the circulating repertoire persistence based on antibody presence/absence, we considered two levels of analysis: first at the sequence level and next at the clonotype level. Here we define persistence as:

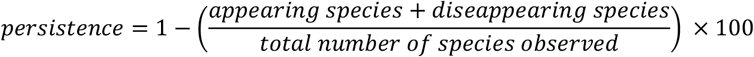

**Figure 2.**
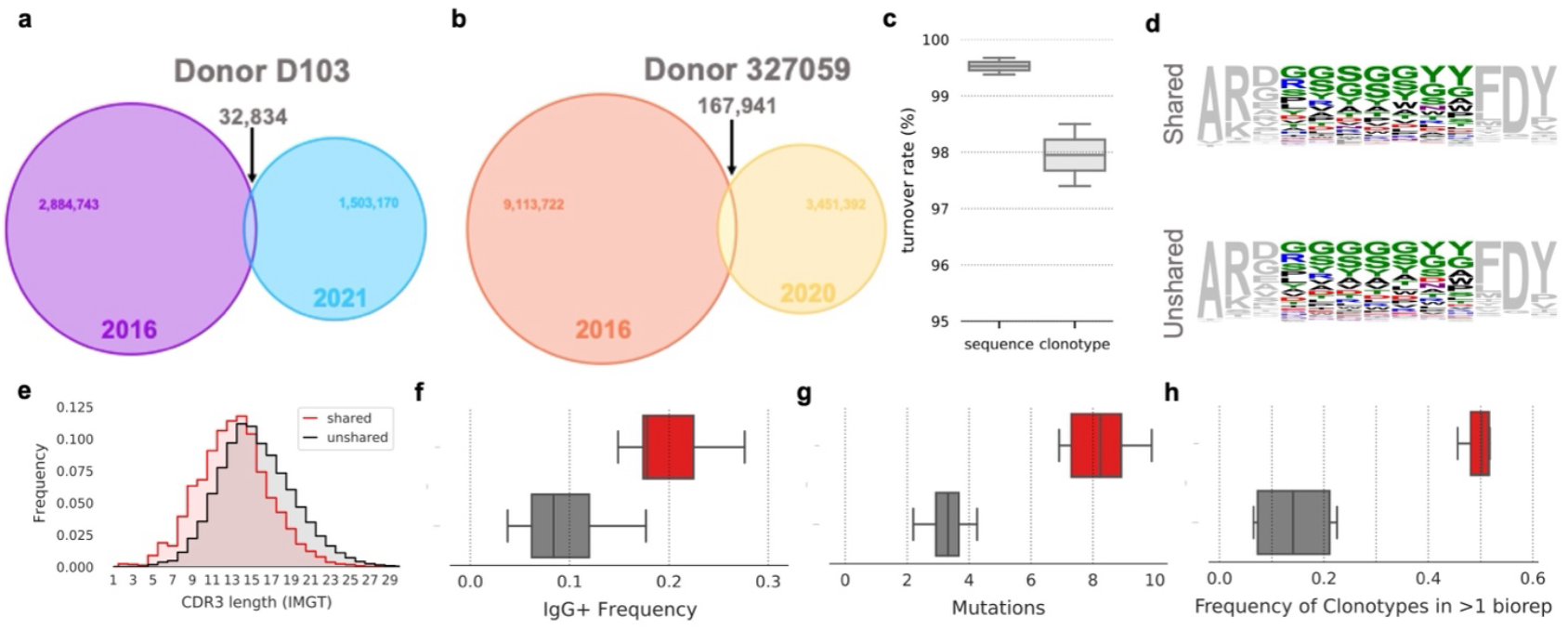
Clonotype and sequence turnover between timepoints. (a) Venn diagram of shared clonotypes between timepoints for subject D103 and (b) for subject 327059. (c) Estimates of total repertoire turnover for sequence and clonotype. (d) Sequence logos of CDRH3s of length 13 amino acids encoded by shared (top) and unshared (bottom) clonotypes between timepoints. (e) Distribution of CDRH3 length for shared and unshared clonotypes between timepoints. (f) Frequency of IgG+ antibodies for shared and unshared clonotypes between timepoints, colored as in (e). Box plots show the median line and span the 25th–75th percentile, with whiskers indicating the 95% confidence interval. (g) Number of V Gene mutations for shared and unshared clonotypes between timepoints, colored as in (e). Box plots show the median line and span the 25th–75th percentile, with whiskers indicating the 95% confidence interval. (h) Frequency of clonotypes found in greater than one biological replicate at a single timepoint for shared and unshared clonotypes between timepoints. Box plots show the median line and span the 25th–75th percentile, with whiskers indicating the 95% confidence interval.

However, this method of estimating persistence is highly sensitive to sampling depth and sequencing bias. Hence, we here only meant to provide tentative results that will require further investigation. Indeed, the present method does not consider varying sampling depth (see Discussion below), resulting in a systematic underestimation of the antibody repertoire persistent fraction. Estimating the true persistence of antibodies and clonotypes over time is a formidable task that would require an extensive statistical modeling effort at minimum, and daunting sequencing capabilities, if only achievable. We calculated the mean clonotype persistence rate over time and obtained a strikingly low value of 2% (2.5% and 1.4% for donors D103 and 327059, respectively). In a similar fashion, persistence rate at the sequence level yielded a mean persistence rate of 0.5% (0.3% and 0.6% for donors D103 and 327959, respectively) (Figure 2c).

Although persistent clonotypes were not distinguishable from clonotypes found only in a single timepoint based solely on the amino acid content of their CDRH3s (Fig 2d), persistent clonotypes were more likely to encode shorter CDRH3s (Fig 2e). Persistent clonotypes found in both timepoints were more frequently of the IgG isotype (Figure 2f) and had a higher mean number of nucleotide V gene mutations (Figure 2g). Our use of multiple biological replicates per sample provides a straightforward means of quantifying repeat occurrence, as any sequences or clonotypes repeatedly observed in different biological replicates must be derived from different cells. Persistent clonotypes were more frequently observed in multiple biological replicates (Figure 2h), suggesting persistent clonotypes to originate from clonally expanded lineages. We cannot exclude the effect of sampling bias on this result, as expanded clonal lineages are more likely to be sampled and thus may be overrepresented among the observed population of persistent clonotypes. However, our analyses of clonotypes shared between separate individuals, which are subject to the same sampling effects as persistent clonotypes within a single donor, do not show an increase in sequences originating from expanded clonal lineages^5^, indicating that sampling alone is not responsible for these phenomena. Taken together, these observations suggest that persistent clonotypes contribute to both repertoire continuity and individuality. The relatively long-lived nature of memory B cells provides a basal level of repertoire continuity that contrasts with the continuous turnover of shorter-lived naïve B cells; however, each individual’s repertoire of persistent clonotypes is the product of their distinct pathogen exposure history and is thus highly personalized. In a sense, the genetic features that define the circulating antibody repertoire (V gene usage, J gene usage, CDRH3 length and amino acid content) are subject to both repertoire shift and drift. Antigen exposure drives episodic repertoire shift, producing substantial and highly individualized changes to the feature distributions of memory B cell repertoires, while naive B cell turnover from cell death and cell replacement causes a more gradual but continuous drift in repertoire composition. Each of these evolutive drivers generates a different signature in the explored features.

### Identification and characterization of persistent public clonotypes

Our previous study quantified the prevalence of repertoire sharing between subjects and found that pairs of subjects shared—on average—0.95% of their respective sampled clonotypes; the number of shared clonotypes between all 10 donors was 0.022%. An examination of public clonotypes between the two subjects at the second timepoint shows public clonotype CDRH3 properties, namely amino acid content and length distributions, are conserved over time, and distinct from non-public clonotypes (Figure S4). We next investigated persistent public clonotypes – those shared between subjects and observed at both timepoints. Remarkably, we found a relatively high number of persistent public clonotypes (Figure 3a). The general CDRH3 amino acid content of persistent public clonotypes was distinct from that of non-public clonotypes (Figure 3b), with shorter CDRH3s more common in persistent as compared to non-persistent public clonotypes (Figure 3c). It is tempting to speculate that these persistent public clonotypes might have a functional role e.g., as convergent Ab responses to common pathogens. Alternatively, the preponderance of Ser and Tyr residues may facilitate the recognition of many antigens^16,17^.

**Figure 3.**
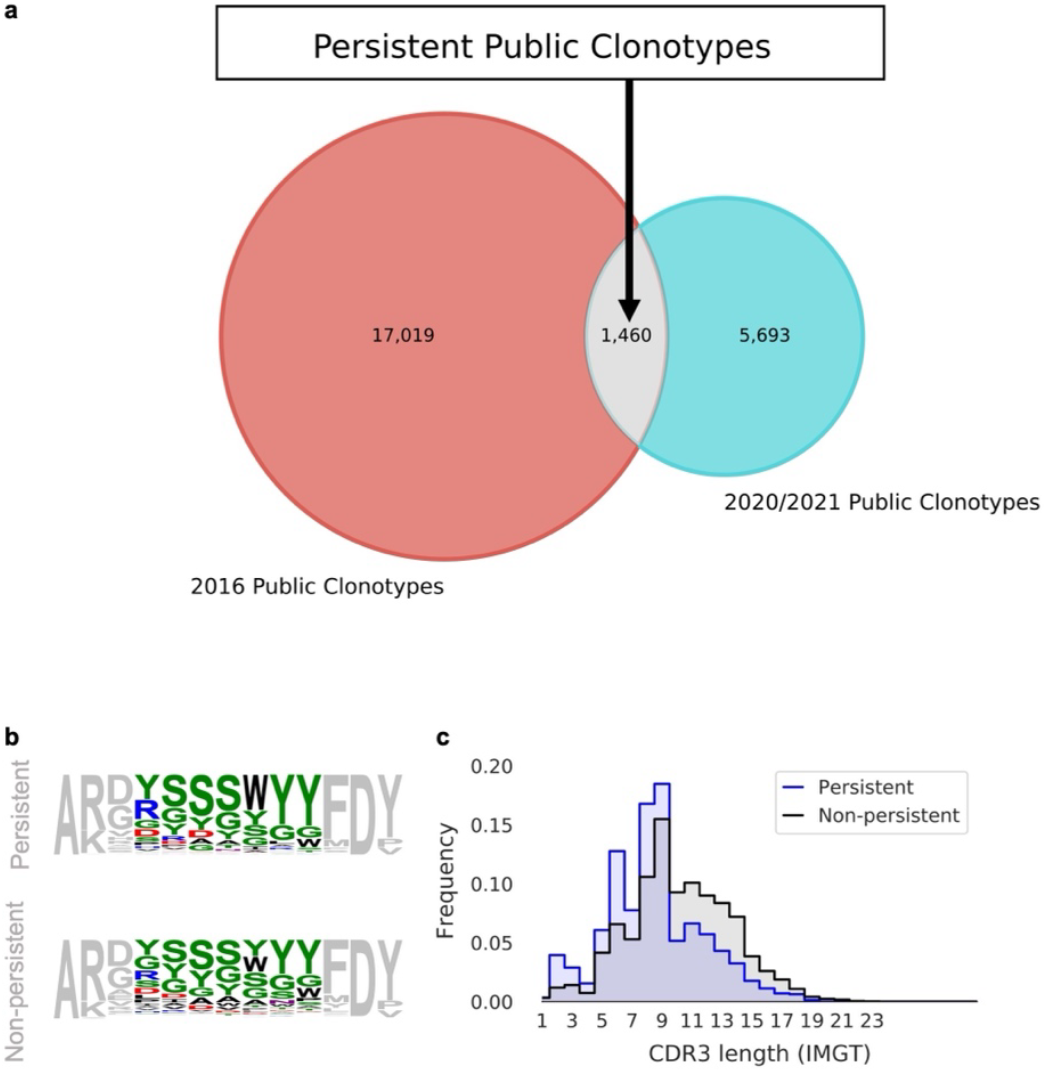
Persistent public clonotypes. (a) Venn diagram of public clonotypes between timepoints. (b) Sequence logos of CDRH3s of length 13 amino acids encoded by persistent (top) and non-persistent (bottom) public clonotypes. (c) Distribution of CDRH3 length for persistent (top) and non-persistent (bottom) public clonotypes.

## DISCUSSION

Overall, our study is consistent with a strikingly low level of conservation in the circulating antibody repertoire over time within a single subject, which is comparable to the expected similarity between unrelated individuals^5^. In other words, the sampled naïve repertoire of an individual 4 years later is suggested to be as different from that sampled from the individual’s prior self as it is between two different individuals at any given time point. This observation, together with the limited number of B cells in any individual, implies the presence of a communal pool of antibody diversity that every individual can access, with the differences between individuals, and within an individual over time, driven primarily by differential sampling of this universal diversity pool. To illustrate this point, one could visualize antibody diversity as a deck of cards^18^ with four distinctive suits - V gene, D gene, J gene, and CDRH3 length. As in a game of chance, everyone is gambling at the immunological casino where they are given four cards at random by the dealer (VDJ recombinase). The hand that any individual player is dealt will vary over time so that the individual will have a unique, time-dependent naïve repertoire to begin countering a given pathogen. That repertoire will be a fraction of the total accessible repertoire because any individual has far less B cells than the size of the accessible repertoire (a healthy adult human has roughly 5 × 10^9^ peripheral blood B cells^12^). This circumstance may disfavor any given individual but will favor the human species because there will always be a fraction of individuals with an advantageous naïve repertoire to build upon to optimally generate immunity to any given threat. In other words, there will always be jackpot winners. The broad result of extensive turnover in the naïve repertoire may have been anticipated given the random nature of recombination to generate the repertoire and the limited half-life of naïve B cells in humans (Macallan et al. 2005).

An important caveat here is the critical limitation of sampling depth in antibody repertoire studies. By necessity, analyses of circulating B cell repertoires can sample only a fraction of the total B cell population, resulting in differences that are the result of insufficient sampling depth but may mimic large-scale repertoire turnover. To mitigate this, we compared multiple biological replicates drawn at each timepoint, each of which are subject to the same sampling biases as samples collected at different timepoints, to establish a baseline level of repertoire difference that can be attributed primarily to sampling bias. Timepoint-matched samples are much more similar than those from different timepoints, meaning the observed temporal shift in repertoire composition must principally be the result of factors other than insufficient sampling. With that said, addressing a bias in human antibody repertoire studies remains a challenging problem. We would propose that animal model studies, where exhaustive sampling of multiple immune compartments is feasible through terminal analyses, may offer more definitive insights into this question.

Our study has also presented an intriguing model of repertoire development, whereby repertoire turnover is strongly influenced by the replacement of circulating naïve B cells from the bone marrow. However, our findings also suggest that immunological exposure and memory formation play a substantial role in preserving circulating clonotypes over time. These factors are especially critical in maintaining higher level repertoire features, as depicted in Figure 1. Therefore, while naïve B cell replacement from the bone marrow may be a strong driver of both repertoire turnover and high-level feature preservation, immunological exposure and memory formation are indispensable for maintaining protective clonotypes. Additionally, we found a relatively high number of persistent public clonotypes. Interestingly, these persistent public clonotypes correspond to shorter CDRH3s, for which two different hypotheses can be formulated. On the one hand, these could correspond to convergent response to shared antigens, such as common infectious diseases or vaccines. The short CDRH3s being here the hallmark of highly specific and mature antibodies. On the other hand, these could be less specific, corresponding to non-specific activation. In this latter case, it would be interesting to know if such persistent public clonotypes with shorter CDRH3s are benefiting the host (providing an overall better baseline to mount a secondary response from) or, quite the opposite, are jeopardizing the antibody response, jamming the immune system with poor quality antibodies arising from the memory compartment. Future studies with larger cohorts, longitudinal follow-up, paired heavy-light chains and multi-tissue sampling will be needed to further characterize the dynamics and regulation of the human antibody repertoire in health and disease. Finally, further research in this area may shed new light on the development of effective vaccines and immunotherapies.

### Limitations of the study

The issue posed by the depth of sampling of antibodies from humans is discussed above. There are further potential limitations to this study that warrant caution when interpreting the results. One issue is the small sample size of the study participants that was dictated by the donors we were able to recruit years after the original study. Variability between individuals could impact the observed differences in both repertoire drift and clonotype usage. Similarly, our study considered participants from the same geographical location, Southern California, potentially influencing the results. Additionally, our study utilized only a limited number of timepoints. Future studies that include larger cohorts, more timepoints and a greater geographical distribution of study participants will be required to corroborate our results.

## MATERIALS AND METHODS

No statistical methods were used to predetermine sample size. The experiments were not randomized, and investigators were not blinded to allocation during experiments and outcome assessment.

### Leukapheresis samples

Full leukopaks (three blood volumes) were obtained from two human subjects (Hemacare). Samples were collected at Hemacare’s Southern California donor center. Sample collection was performed under a protocol approved by the Institutional Research Boards of Scripps Research and Hemacare. Informed consent was obtained from each subject. Both subjects were healthy, HIV-negative adults between the ages of 18 and 30 with no reported acute illness in the 14 days before leukapheresis. Immediately upon receipt of the leukopak, peripheral blood mononuclear cells were purified by gradient centrifugation and cryo-preserved.

### Amplification strategy and primer bias

We elected to use RNA as the template for antibody variable gene amplification, as this focuses our analysis on productive heavy-chain rearrangements and permits the use of amplification primers that anneal to the CH1 region (owing to the presence of an intron between the JH gene and CH region, the use of CH1 primers is not feasible when amplifying from DNA). The decision to use RNA has some inherent downsides, however—primarily the likelihood of overrepresentation of transcriptionally active B cells (namely, memory B cells and plasmablasts). It should be noted that the use of molecular barcodes, which enable identification and collapsing of reads that originate from the same RNA molecule, will not correct this problem. To reduce the influence of multiplexed primer sets on the resulting composition of antibody genes that are amplified, we designed an amplification strategy that limits the use of multiplexed primers that anneal to the V-gene region in an attempt to reduce primer bias during amplification. Following cDNA synthesis, second-strand synthesis was performed using multiplexed V-gene primers that encode an overhang that comprises a portion of the Illumina adapters required for next-generation sequencing. V-gene primers were then enzymatically removed before subsequent amplification of the antibody genes using the conserved overhang as the primer annealing site. Thus, the multiplexed V-gene primers were only used for a single round of amplification.

### Antibody gene amplification

For each subject, total RNA was separately isolated from 6 aliquots of approximately 4 × 10^8^ cryo-preserved peripheral blood mononuclear cells (RNeasy Maxi, Qiagen). For each RNA aliquot, antibody genes were amplified in triplicate (18 total samples per subject), with each of the technical replicates processed independently and starting with a separate aliquot of the RNA sample. To minimize the likelihood of cross contamination between subjects, reverse transcription and PCR reactions for each subject were processed in isolation, such that samples from two different subjects were never in proximity during amplification reaction preparation. All primers are listed in Briney et al., 2019^5^. To increase the sequencer-perceived nucleotide diversity during each sequencing cycle, ‘offsets’ were added to the reverse-transcription and second-strand synthesis primers. Three sets of these primers were synthesized, with each set containing 2, 4 or 6 random nucleotides at the offset position. These offsets stagger the conserved constant and framework regions and result in much higher diversity during each sequencing cycle and minimize the required PhiX spike. cDNA synthesis was performed on 11 μl of RNA using 10 pmol of each primer in a 20-μl total reaction (SuperScript III, Thermo Fisher Scientific), using the manufacturer’s protocol and the following thermal cycling program: 55 °C for 60 min, 70 °C for 15 min. Residual primers and dNTPs were degraded enzymatically (ExoSAP-IT, Thermo Fisher Scientific), according to the manufacturer’s protocol. The entire enzyme-treated cDNA synthesis product was used in a 100-μl second-strand synthesis reaction using 10 pmol of each primer (HotStarTaq Plus, Qiagen) using the following thermal cycling protocol: 95 °C for 5 min, 55 °C for 30 s, 72 °C for 10 min. Residual primers and dNTPs were again degraded enzymatically (ExoSAP-IT) and dsDNA was purified using 0.8 volumes of SPRI beads (AmpureXP, Beckman Coulter Genomics) and eluted in 50 μl of water. Antibody genes were amplified using 40 μl of eluted dsDNA and 10 pmol of each primer in a 100-μl total reaction volume (HotStarTaq Plus), using the following thermal cycling program: 95 °C for 5 min; 25 cycles of: 95 °C for 30 s, 58 °C for 30 s, 72 °C for 2 min; 72 °C for 10 min. DNA was purified from the PCR reaction product using 0.8 volumes of SPRI beads (AmpureXP) and eluted in 50 μl of water. Subsequently, 10 μl of the eluted PCR product was used in a final indexing PCR (HotStarTaq Plus) using 10 pmol of each primer in 100-μl total reaction volume and using the following thermal cycling program: 95 °C for 5 min; 10 cycles of: 95 °C for 30 s, 58 °C for 30 s, 72 °C for 2 min; 72 °C for 10 min. PCR products were purified with 0.7 volumes of SPRI beads (SPRIselect, Beckman Coulter Genomics) and the entire set of samples from a single subject was eluted in a single 120-μl volume of water.

### Sequencing

SPRI-purified sequencing libraries were initially quantified using fluorometry (Qubit, Thermo Fisher Scientific) before size determination using a Bioanalyzer (Agilent 2100). Libraries were re-quantified using qPCR (KAPA Biosystems) before sequencing on an Illumina NovaSeq 6000 using 2 × 250-bp SP chemistry.

### Processing of raw sequencing data

Raw paired FASTQ files were quality checked with FASTQC (www.bioinformatics.babraham.ac.uk/projects/fastqc/). Because the 5′ end of each paired read encodes the unique molecular identifier (UMI), reads were quality trimmed only at the 3′ end using Sickle^19^ (www.github.com/najoshi/sickle), using a window size of 0.1 times the length of the read, minimum average window quality score of 20, and a minimum read length after trimming of 50 nucleotides. Because UMIs are located on the ‘outside’ of the gene-specific primers used for amplification, primer trimming was delayed until after UMI processing. Processed reads were quality checked again using FASTQC, and paired reads were merged with PANDAseq using the default (simple_Bayesian) merging algorithm^20^.

### Molecular barcodes

Although sequencing libraries were constructed to encode molecular barcodes on both ends of the amplicon, we observed low-level PCR recombination that produced ‘barcode swapping’, causing the frequency of these amplification artefacts to be amplified. In essence, a partial amplification product—composed of a CDRH3 and an incomplete VH gene—was able to prime a different antibody sequence and continue amplification, producing a hybrid VH gene. This hybrid amplicon encodes the 3′ molecular barcode from the primary antibody recombination and the 5′ molecular barcode from the second. The barcode swapping creates a unique barcode pair, forcing the hybrid sequence to be binned and processed separately. To minimize the effects of such barcode swapping, we binned sequences using only the 3′ molecular barcode. Because the likelihood of UMI collisions was relatively high given the sequencing depth, the CDRH3 nucleotide sequences of each UMI bin containing more than one sequence were clustered at high identity (90%) and a consensus sequence was computed for each cluster. For UMI bins containing only a single sequence, the lone sequence was used as the representative for the respective UMI bin. The majority of UMI bins contained only a single sequencing read. As such, the UMIs were not used primarily for error correction, but as a means for correcting differential representation arising from stochastic or primer-driven amplification biases. Mutation frequencies in the IgM and IgG sequence populations provide empirical evidence of a low amplification and sequencing error rate that corroborates sequencer-derived quality metrics.

### Germline-gene assignment and annotation

Adapters and V-gene amplification primers (used for second-strand synthesis) were removed using cutadapt^21^. cDNA synthesis primers, which anneal to the CH1 region, were not removed because this region is needed to determine the isotype. Sequences were annotated with abstar^10^and two output formats were generated: a comprehensive JSON-formatted output, which was imported into a MongoDB database; and a tabular CSV-formatted output, which is suitable for direct parsing or conversion to Parquet for querying on a Spark cluster.

### Morisita–Horn similarity

Antibody sequences from each subject were reduced to only the V-gene, J-gene and CDRH3 length (and were randomly subsampled with replacement at sample sizes ranging from 10^1^ to 10^7^). The frequency of each V-gene, J-gene and CDRH3 length was computed, and the frequency distributions from two donors were used to compute the Morisita–Horn similarity index:

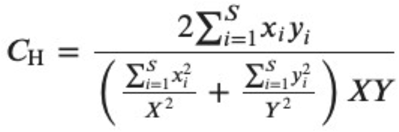

in which x_*i*_ is the number of times the V-gene, J-gene and CDRH3 length *i* is represented in one sample of size *X*; and y_*i*_ is the number of times the V-gene, J-gene and CDRH3 length *i* is represented in a second sample of size *Y*.

### Classification of repertoires by subject and timepoint

Repertoires were classified using a one-versus-rest support-vector-machine classifier. Classifier training and evaluation were performed in Python using the scikit-learn framework. It is important to note that this classification was performed using only 4 samples and expanding the subject and timepoint pool to thousands or millions, while maintaining classifier accuracy, would likely require much larger training datasets and/or the inclusion of additional sequence features to supplement the V-gene, J-gene and CDRH3 length.

### Turnover Calculation

Antibody sequences from each subject were reduced to unique sequences and unique clonotypes, and turnover was calculated for both sequences and clonotypes separately as defined^15^: (Number of appearing species + number of disappearing species) / (total number of species observed) x 100.

### Statistical Calculations

Statistical calculations were performed in Python using SciPy^22^ and Seaborn^23^.

## Code availability

Code for in this manuscript is available at github.com/CollinJ0/grp2_paper. Abstar is available at github.com/brineylab/abstar. Code for molecular barcode processing is available at www.github.com/brineylab/abtools.

## Data availability

Sequence data that support the findings in this study are available at the NCBI Sequencing Read Archive under BioProject numbers PRJNA406949 and PRJNA1005308.

## Acknowledgments

The authors thank the study subjects for their participation.

## AUTHOR CONTRIBUTIONS

Conceptualization: CJ, DRB, BB

Data generation and analysis: CJ, BN, RB

Manuscript preparation and revisions: CJ, BN, RB, BB, DRB

## FUNDING

This work was supported by the National Institute of Allergy and Infectious Diseases UM1AI144462 (DRB and BB), U19AI135995 (BB), P01-AI177683 (BB), R01-AI171438 (BB),

P30-AI036214 (BB); the International AIDS Vaccine Initiative through the Neutralizing Antibody Consortium, SFP1849 (DRB); the Ragon Institute of MGH, MIT and Harvard (DRB); and the Pendleton Foundation.

## DECLARATION OF INTERESTS

BB is an equity shareholder in Infinimmune and a member of their Scientific Advisory Board.

## Notes

https://www.github.com/CollinJ0/grp2_paper

