## Supplementary Information for "Longitudinal analysis of drift in the circulating human antibody repertoire over four years"

**This PDF file includes:**

Figures S1 to S4 and Table S1

**Supplemental Figure 1. V/J frequency correlations of technical and biological replicates.** For each sample, the frequency of V and J combinations was compared for technical replicates (left panels) or biological replicates (right panels). The coefficient of determination (r2) is shown for each plot.

### Figure S1

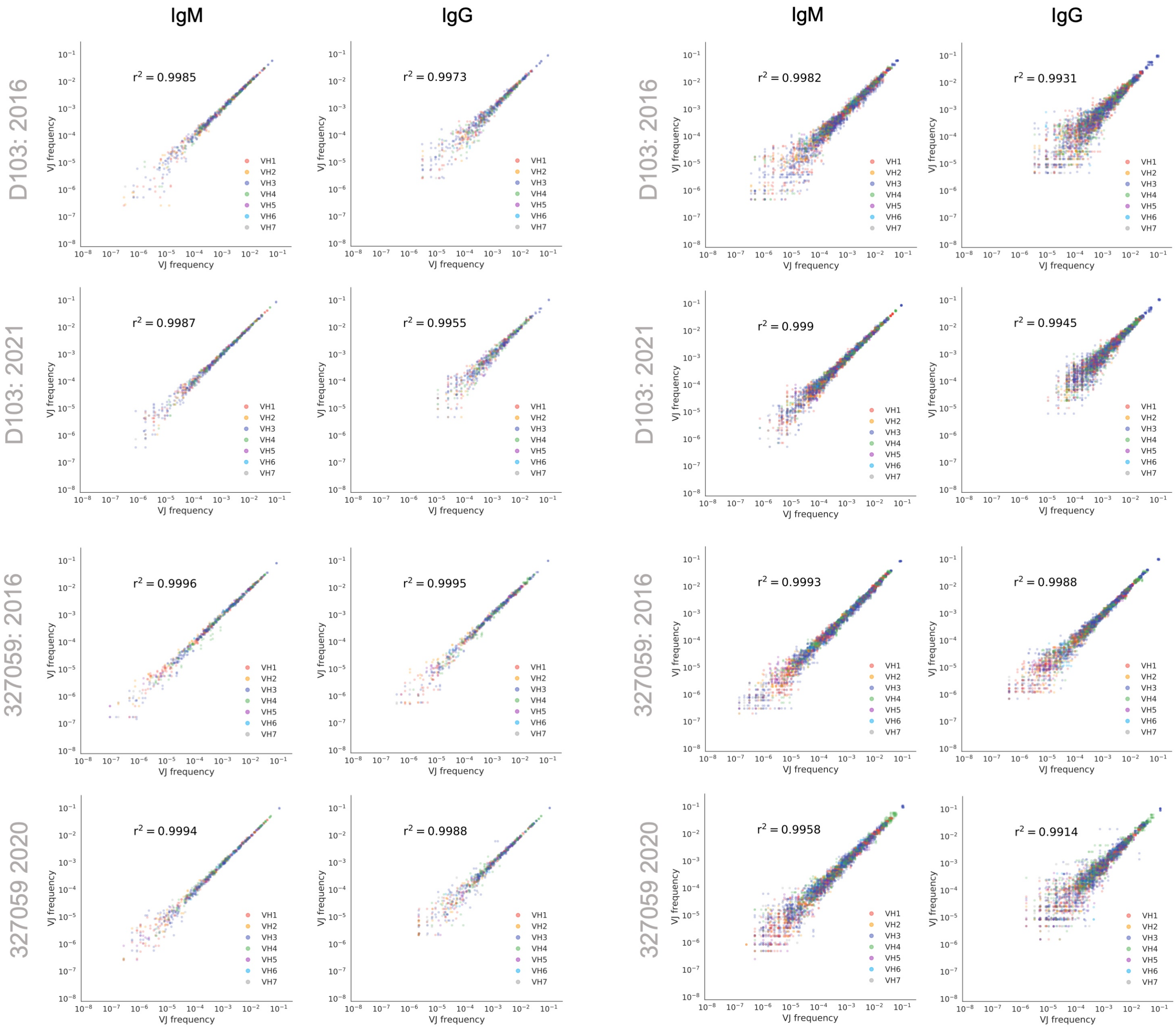
IgM
IgG
IgM
IgG
D103: 2016
D103: 2021
327059: 2016
327059 2020
r$^2$ = 0.9985
r$^2$ = 0.9973
r$^2$ = 0.9982
r$^2$ = 0.9931
r$^2$ = 0.9987
r$^2$ = 0.9955
r$^2$ = 0.999
r$^2$ = 0.9945
r$^2$ = 0.9996
r$^2$ = 0.9995
r$^2$ = 0.9993
r$^2$ = 0.9988
r$^2$ = 0.9994
r$^2$ = 0.9988
r$^2$ = 0.9958
r$^2$ = 0.9914
VJ frequency
VH1
VH2
VH3
VH4
VH5
VH6
VH7

**Supplemental Figure 2. V/J frequency correlations between timepoints.** For each subject, the frequency of V and J combinations was compared between timepoints for all isotypes (left), IgM sequence (center) and IgG sequences (right). The coefficient of determination (r2) is shown for each plot.

Figure S2

### Inter-year comparisons

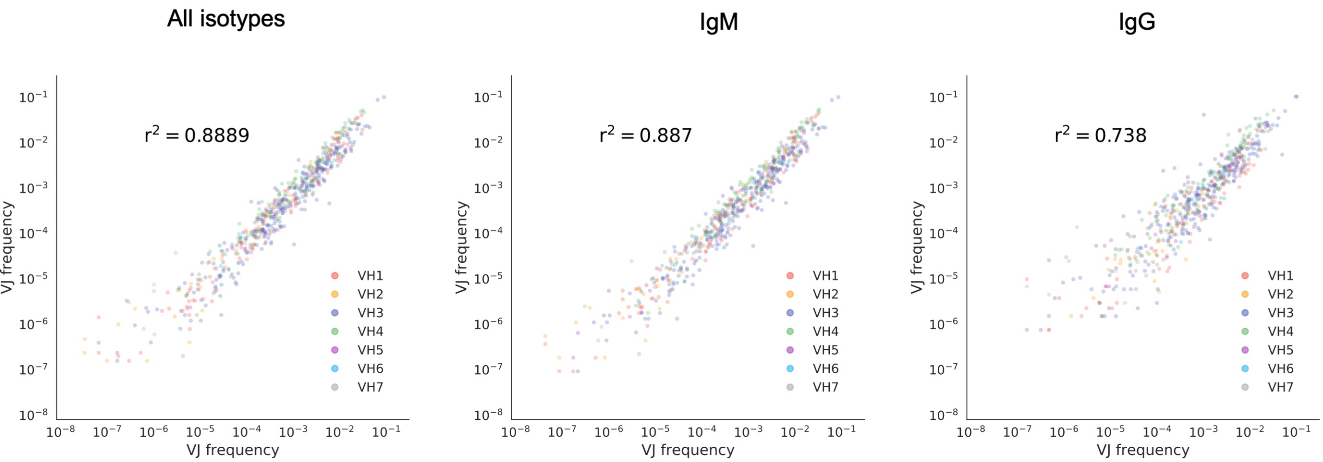

**Supplemental Figure 3. Percent shared clonotypes (solid line) and sequences (dotted line) verses sample depth.** Lines indicate mean frequency of 10 bootstrap samplings.

#### Figure S3

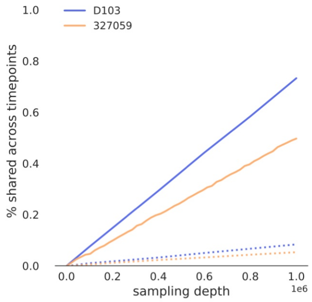
D103
327059
% shared across timepoints
1.0
0.8
0.6
0.4
0.2
0.0
0.0
0.2
0.4
0.6
0.8
1.0
1e6
sampling depth

**Supplemental Figure 4. Public clonotype properties for each timepoint.** Top panel shows sequence logos of CDRH3s of length 13 amino acids encoded by shared (top) and unshared (bottom) clonotypes between donors at each timepoint. Bottom panel shows distribution of CDRH3 length for shared and unshared public clonotypes between donors at each timepoint.

Figure S4

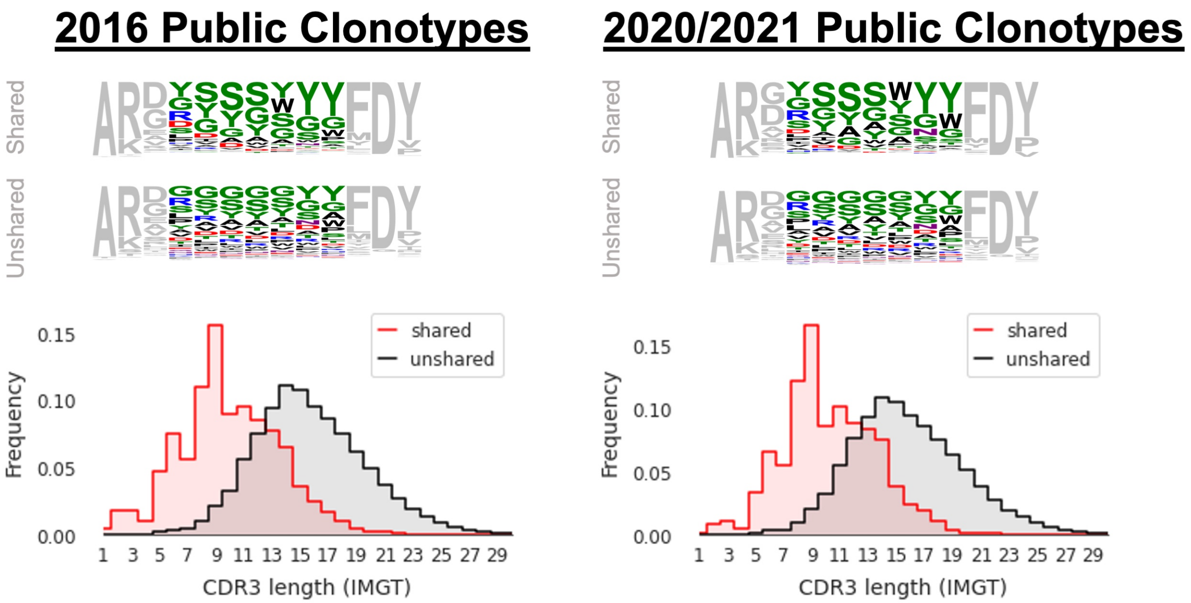

2016 Public Clonotypes
2020/2021 Public Clonotypes
Shared
Unshared
Shared
Unshared
Frequency
shared
unshared
CDR3 length (IMGT)
Frequency
shared
unshared
CDR3 length (IMGT)

| Subject | Timepoint #1 | Timepoint #2 | Age at Timepoint #1 | Age at Timepoint #2 | Raw reads | Consensus sequences | Clonotypes |
|---|---|---|---|---|---|---|---|
| 327059 | 11/22/16 | 12/2/20 | 26 | 30 | 801,514,273 | 44,035,482 | 12,900,996 |
| D103 | 11/17/16 | 4/1/21 | 25 | 30 | 473,638,628 | 16,862,355 | 4,453,581 |

**Supplemental Table 1. Per-subject timepoint information and sequencing statistics.**
